# MERS-CoV endoribonuclease and accessory proteins jointly evade host innate immunity during infection of lung and nasal epithelial cells

**DOI:** 10.1101/2021.12.20.473564

**Authors:** Courtney E. Comar, Clayton J. Otter, Jessica Pfannenstiel, Ethan Doerger, David M. Renner, Li Hui Tan, Stanley Perlman, Noam A. Cohen, Anthony R. Fehr, Susan R. Weiss

## Abstract

Middle East respiratory syndrome coronavirus (MERS-CoV) emerged into humans in 2012, causing highly lethal respiratory disease. The severity of disease may be in part because MERS-CoV is adept at antagonizing early innate immune pathways – interferon (IFN) production and signaling, protein kinase R (PKR), and oligoadenylate synthetase ribonuclease L (OAS/RNase L) – generated in response to viral double-stranded (ds)RNA generated during genome replication. This is in contrast to SARS-CoV-2, which we recently reported activates PKR and RNase L and to some extent, IFN signaling. We previously found that MERS-CoV accessory proteins NS4a (dsRNA binding protein) and NS4b (phosphodiesterase) could weakly suppress these pathways, but ablation of each had minimal effect on virus replication. Here we investigated the antagonist effects of the conserved coronavirus endoribonuclease (EndoU), in combination with NS4a or NS4b. Inactivation of EndoU catalytic activity alone in a recombinant MERS-CoV caused little if any effect on activation of the innate immune pathways during infection. However, infection with recombinant viruses containing combined mutations with inactivation of EndoU and deletion of NS4a or inactivation of the NS4b phosphodiesterase promoted robust activation of the dsRNA-induced innate immune pathways. This resulted in ten-fold attenuation of replication in human lung derived A549 and primary nasal cells. Furthermore, replication of these recombinant viruses could be rescued to the level of WT MERS-CoV by knockout of host immune mediators MAVS, PKR, or RNase L. Thus, EndoU and accessory proteins NS4a and NS4b together suppress dsRNA-induced innate immunity during MERS-CoV infection in order to optimize viral replication.

**Importance:** Middle East Respiratory Syndrome Coronavirus (MERS-CoV) causes highly lethal respiratory disease. MERS-CoV encodes several innate immune antagonists, accessory proteins NS4a and NS4b unique to the merbeco lineage and the nsp15 protein endoribonuclease (EndoU), conserved among all coronaviruses. While mutation of each antagonist protein alone has little effect on innate immunity, infections with recombinant MERS-CoVs with mutations of EndoU in combination with either NS4a or NS4b, activate innate signaling pathways and are attenuated for replication. Our data indicate that EndoU and accessory proteins NS4a and NS4b together suppress innate immunity during MERS-CoV infection, to optimize viral replication. This is in contrast to SARS-CoV-2 which activates these pathways and consistent with greater mortality observed during MERS-CoV infection compared to SARS-CoV-2.

## Introduction

Middle East respiratory syndrome coronavirus (MERS-CoV) first emerged in 2012, in Saudi Arabia (1), and was the second of three zoonotic coronaviruses to emerge into humans in the 21^st^ century, following severe acute respiratory syndrome coronavirus (SARS-CoV) in 2002 and preceding SARS-CoV-2 in 2019. MERS-CoV is a highly pathogenic betacoronavirus, of the merbeco lineage, that has caused 888 deaths in 2578 laboratory confirmed cases (as of November 2021) according to the World Health Organization, amounting to a case-fatality rate of over 34%. MERS-CoV circulates in its natural reservoir, dromedary camels, and closely related viruses have been found in bats, suggesting that MERS-CoV descended from a bat virus (2–7).

Coronavirus genomes are large positive sense single-stranded (ss)RNA with a 5’ cap and 3’ polyA tail. Specifically, the MERS-CoV genome is 30,119 nucleotides in length. The 5’ two-thirds of the coronavirus genome encodes the conserved replicase proteins in Orf1a and Orf1b, involved in several processes such as RNA replication and transcription, RNA capping and processing, proteolytic processing of replicase proteins, and host innate immune antagonism. The 3’ one-third of the genome encodes the structural genes and the unique, lineage-specific accessory genes, which are not necessary for viral replication but play important roles in immune evasion and viral pathogenesis.

Early in infection, coronaviruses establish transcription/replication complexes within the endoplasmic reticulum, associated double membrane vesicles. Replication of the coronavirus genome RNA and transcription of mRNAs takes place in these complexes and generates dsRNA intermediates. This dsRNA provides a pathogen associate molecular pattern (PAMP) that is detected by several different host pattern recognition receptors (PRRs), leading to activation of innate immune antiviral pathways. Melanoma differentiation-associated protein 5 (MDA5) is a PRR that detects coronavirus dsRNA (8) and induces signaling through mitochondrial antiviral-signaling protein (MAVS), which leads to activation of interferon regulatory factors (IRF), and transcription of type I and III interferons (IFN) (9). IFNs are produced and secreted by the infected cell and can signal in autocrine or paracrine fashion. Downstream JAK/STAT signaling and phosphorylation leads to expression of several hundred interferon stimulatory genes (ISGs) which produce an antiviral state. Protein Kinase R (PKR) is another cytoplasmic PRR activated by dsRNA. Upon sensing dsRNA, PKR autophosphorylates and then phosphorylates translation initiation factor eIF2α leading to translation arrest (10). Oligoadenylate synthetases (OAS1-3), upon detecting dsRNA, produce 2’,5’ -oligoadenylates (2-5A) that activate the host enzyme ribonuclease L (RNase L) which cleaves viral and host single-stranded (ss)RNA (11, 12). Activation of each these pathways leads to restriction of virus replication and spread. RNase L and PKR activation can promote IFN production, cellular stress, inflammation, and/or apoptotic death (13–16). Many viruses, including coronaviruses, have evolved various mechanisms for counteracting these pathways. Importantly, each of the three pathways can be activated independently of the others (17–19); therefore, if IFN signaling is inhibited during infection as is the case for some viruses, the OAS/RNase L or PKR pathways can still be activated. In addition, since OASs and PKR are ISGs, the PKR and OAS/RNase L pathways can be further upregulated by IFN.

Coronavirus accessory proteins contribute to differences between the three lineages of betacoronaviruses. The merbeco virus MERS-CoV is particularly efficient at shutting down host innate immune pathways, including IFN signaling, PKR, and OAS/RNase L, while the sarbecovirus, SARS-CoV-2 activates PKR and RNase L and weakly activates IFNs (18). Two accessory proteins encoded by MERS-CoV have been shown to play important roles in this innate immune antagonism (20–22). NS4a and NS4b are both encoded on mRNA4 by ORFs 4a and 4b respectively. NS4a is a dsRNA binding protein and as such has been reported to block IFN induction and PKR activation (21, 23–27). NS4b has at least two functional domains, a phosphodiesterase (PDE) and an amino terminal nuclear localization signal (NLS). The PDE degrades 2-5A preventing activation of RNase L and weakly inhibits IFN production (21, 22). The amino terminal nuclear localization signal (NLS) confers expression mostly in the nucleus and has been reported to block NFκB translocation to the nucleus (22, 28–31). Previously we found that a recombinant mutant MERS-CoV ablated for NS4a expression induced low levels of phosphorylated PKR and no detectable phosphorylation of eIF2α, as well as a low level of type I and III IFN expression (21), and a recombinant MERS-CoV with an inactivated NS4b PDE led to mild activation of RNase L and IFN induction (21, 22). Notably, neither the embecoviruses (for example human coronavirus (HCoV)-OC43 or murine coronavirus (MHV)) nor sarbeco lineages (for example SARS-CoV-2) encode a dsRNA binding protein. While the sarbecoviruses do not express a PDE, the embecoviruses do encode a PDE in the NS2 protein. However, in contrast to NS4b, this PDE lacks an NLS and thus is expressed completely in the cytoplasm (32). These differences in accessory proteins between CoVs likely contribute to differences in pathogenesis and immune evasion.

Nsp15 is a conserved coronavirus replicase protein encoded in Orf1b that contains an endoribonuclease (EndoU) domain with two catalytic histidine residues. Murine coronavirus (MHV) EndoU has been shown to limit dsRNA accumulation and consequently reduces IFN production and signaling as well as activation of OAS/RNase L and PKR pathways during infection of murine bone marrow-derived macrophages (33, 34). While the exact mechanism of action of EndoU is not completely understood, two different mechanisms have been reported. One report concludes that EndoU cleaves genomic RNA to limit the production of dsRNA (35) and the other that EndoU degrades polyU from the 5’ end of negative strand RNA, eliminating a PAMP for sensors (36). The role of MERS-CoV EndoU activity in antagonizing innate immunity has yet to be examined during infection of human cells.

Using a group of recombinant mutant viruses, we investigated the role of EndoU, alone and in combination with NS4a or NS4b in innate immune antagonism during MERS-CoV infection. In addition to investigating immune evasion by MERS-CoV in the lung-derived cell line A549^DPP4^, which expresses MERS-CoV receptor dipeptidyl peptidase 4, we extended the study of these host-pathogen immune interactions to primary patient-derived nasal epithelial cells in air-liquid interface cultures. Our findings indicate that MERS-CoV EndoU together with its accessory proteins effectively shut down innate immune pathways to optimize replication.

## Results

### Construction of MERS-CoV recombinant viruses and replication kinetics in VeroCCL81 and A549^DPP4^ cells

In order to study the effects of EndoU activity on the dsRNA-induced antiviral innate immune pathways during MERS-CoV infection, several recombinant mutants were constructed using the Lambda Red recombination system and a Bacterial Artificial Chromosome (BAC) that encodes the full-length MERS-CoV genome (37). Recombinant viruses (summarized in Fig 1A) were MERS-CoV-nsp15^H231A^ with an amino acid substitution inactivating EndoU, MERS-CoV-ΔNS4a with interruption of NS4a expression by insertion of two termination codons at positions 11 and 12, MERS-CoV-NS4b^H182R^ with an amino acid substitution inactivating the PDE, and double mutants MERS-CoV-nsp15^H231A^/ΔNS4a and MERS-CoV-nsp15^H231A^/ NS4b^H182R^. We confirmed that MERS-ΔNS4a and -nsp15^H231A^/ΔNS4a infection produced undetectable expression of NS4a protein by Western blot (Fig 1B). As we observed previously, an NS4b^H182R^ mutant expressed less NS4b than WT MERS-CoV (Fig 1B) (21).

**Figure 1:**
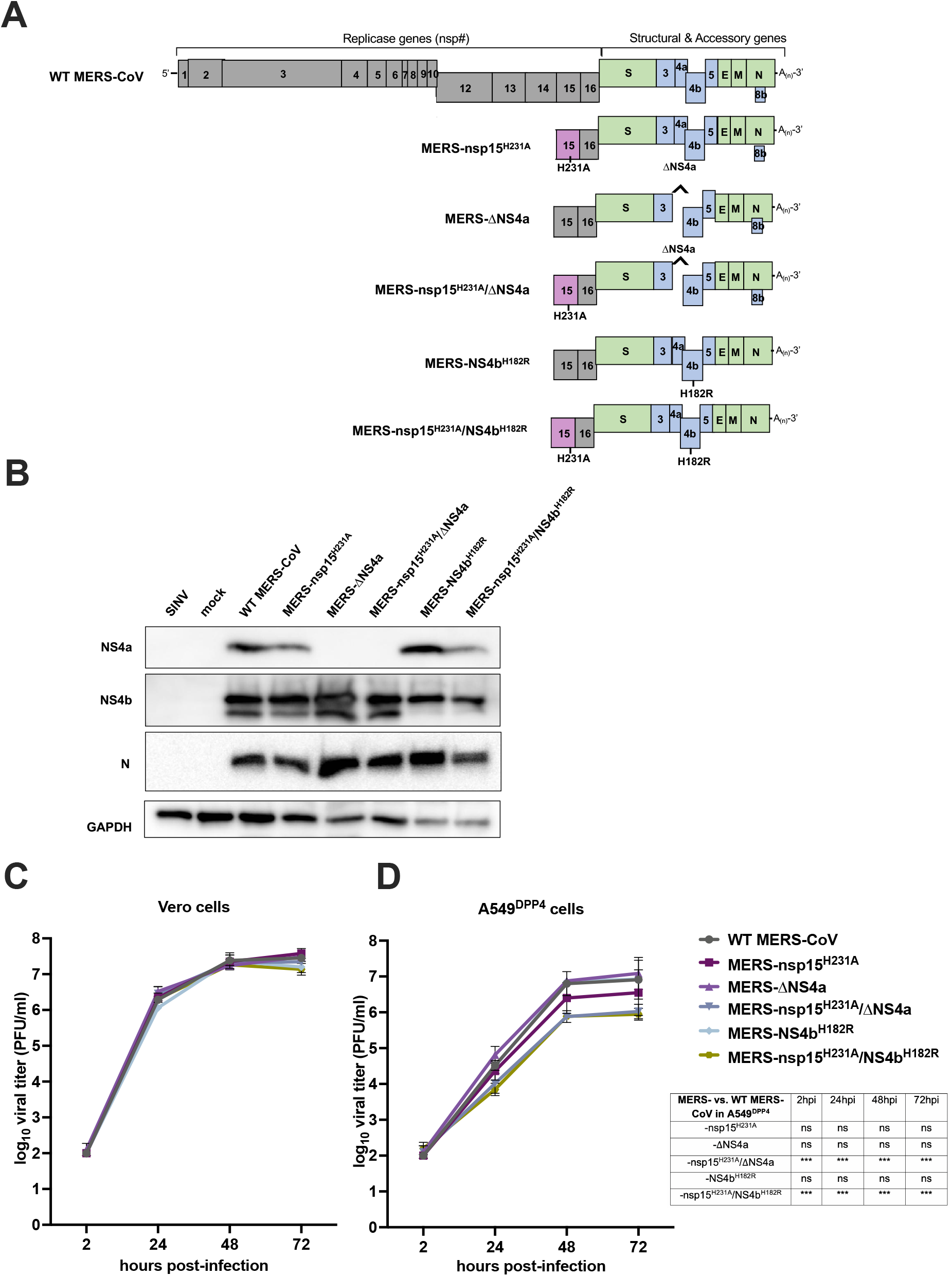
Recombinant MERS-CoV design and replication kinetics. (A) Diagram of the MERS-CoV genome including the replicase locus [encoding 16 nonstructural proteins (nsp)], structural genes and accessory genes is shown along with diagram of each recombinant mutant constructed using the BAC reverse genetics system. (B) A549^DPP4^ cells were infected at MOI=5 with the indicated viruses and protein lysates harvested at 24hpi. Expression of NS4a and NS4b was determined by SDS-PAGE and Western blot. (C) Vero cells were infected in triplicate at an MOI of 1 with the indicated viruses. Supernatants were collected at the indicated hours post-infection (hpi), and infectious virus was quantified by plaque assay. Data are displayed as means ± standard deviation (SD). None of the data were statistically significant by repeated measures two-way ANOVA (D) A549^DPP4^ cells were infected in triplicate with the indicated viruses at an MOI of 1 and replication was quantified as in (C). Data are displayed as means ± standard deviation (SD). Statistical significance of each recombinant virus compared to WT MERS-CoV was calculated by repeated measures two-way ANOVA: *, *P*≤0.05; **, *P* ≤ 0.01; ***, *P* ≤ 0.001; ****, *P* ≤ 0.0001. Data that were not statistically significant are labeled ns. Data from (B,C) are from one representative of three independent experiments.

We compared replication of the mutant viruses with WT MERS-CoV initially in African Green Monkey kidney Vero CCL81 (Vero) cells, defective in IFN signaling. Cells were infected at MOI=1 and supernatant samples harvested at 2, 24, 48, and 72 hours-post infection (hpi). Quantification of infectious virus was completed by standard viral plaque assay on Vero cells as previously described (18). Mutant viruses replicated to similar levels as WT MERS-CoV in three independent experiments (Fig 1C), indicating the absence of inherent replication defects. To investigate the effects of dsRNA-induced innate immune activity on replication of these viruses, A549^DPP4^ cells (with intact IFN, PKR, and RNase L pathways) were infected at MOI=1 and infectious virus quantified as above. At 48 and 72 hpi, we observed an approximate 1 log_10_ PFU/mL decrease in infectious virus produced in the MERS-nsp15^H231A^/ΔNS4a and - nsp15^H231A^/NS4b^H182R^ mutants compared to WT MERS-CoV that reached statistical significance (Fig 1D). Interestingly, mutation of the EndoU catalytic site alone, MERS-nsp15^H231A^, resulted in a small but not statistically significant defect in infectious virus production. No differences were noted in MERS-ΔNS4a or -NS4b^H182R^ replication. This is consistent with our previous finding of only mild effects on viral replication in a mutant lacking expression of either of these two accessory proteins (21).

### DsRNA accumulation is increased when EndoU is inactive during MERS-CoV infection

Next, we sought to examine production of dsRNA during infection with MERS-CoV recombinants containing catalytic inactivation mutations in EndoU (nsp15^H231A^) compared with WT MERS-CoV. Detection of dsRNA was assessed by immunofluorescence assay (IF) using the monoclonal antibody J2 directed against dsRNA and imaged using widefield microscopy. In several experiments we observed increased expression of dsRNA when EndoU was inactivated, including during infection with MERS-nsp15^H231A^ and -nsp15^H231A^/ΔNS4a compared to WT MERS-CoV (Fig 2A), consistent with previous findings for inactivation of MHV EndoU (33, 36). To quantify this observation and to extend it to the other MERS-CoV mutants, we infected A549^DPP4^ at MOI=5 on glass coverslips and fixed with 4% paraformaldehyde. We performed combined fluorescent *in situ* hybridization (FISH)/IF using J2 antibody to detect dsRNA, antiserum against the viral primase, nsp8, a component of the viral polymerase complex and therefore a marker for virus replication/transcription complexes (38), and oligonucleotide probes to detect nucleocapsid mRNA to define the cytoplasm. We observed brighter staining of dsRNA in MERS-nsp15^H231A^, -nsp15^H231A^/ΔNS4a, and -nsp15^H231A^/NS4b^H182R^ compared to WT MERS-CoV at 48hpi (Fig 2B). Fiji software was used for quantification, as described in detail in the Materials and Methods section. Briefly, due to extensive cell-to-cell fusion (syncytia), we were unable to outline individual infected cells for quantification. To quantify dsRNA, we outlined infected areas of images (syncytia and single cells) using nucleocapsid RNA staining to define the cytoplasm and called them regions of interest (ROIs). We measured the mean gray value (MGV) of fluorescence signal of dsRNA or nsp8 within each ROI and recorded the ratio of dsRNA MGV over nsp8 MGV for each ROI. These ratios were compared across infection with mutant and WT viruses at 48hpi in A549^DPP4^ cells (Fig 2C). We observed statistically significant increases in dsRNA/nsp8 ratios in MERS-nsp15^H231A^, -nsp15^H231A^/ΔNS4a, and - nsp15^H231A^/NS4b^H182R^ compared to WT MERS-CoV but not MERS-ΔNS4a or -NS4b^H182R^ at 48hpi.

**Figure 2:**
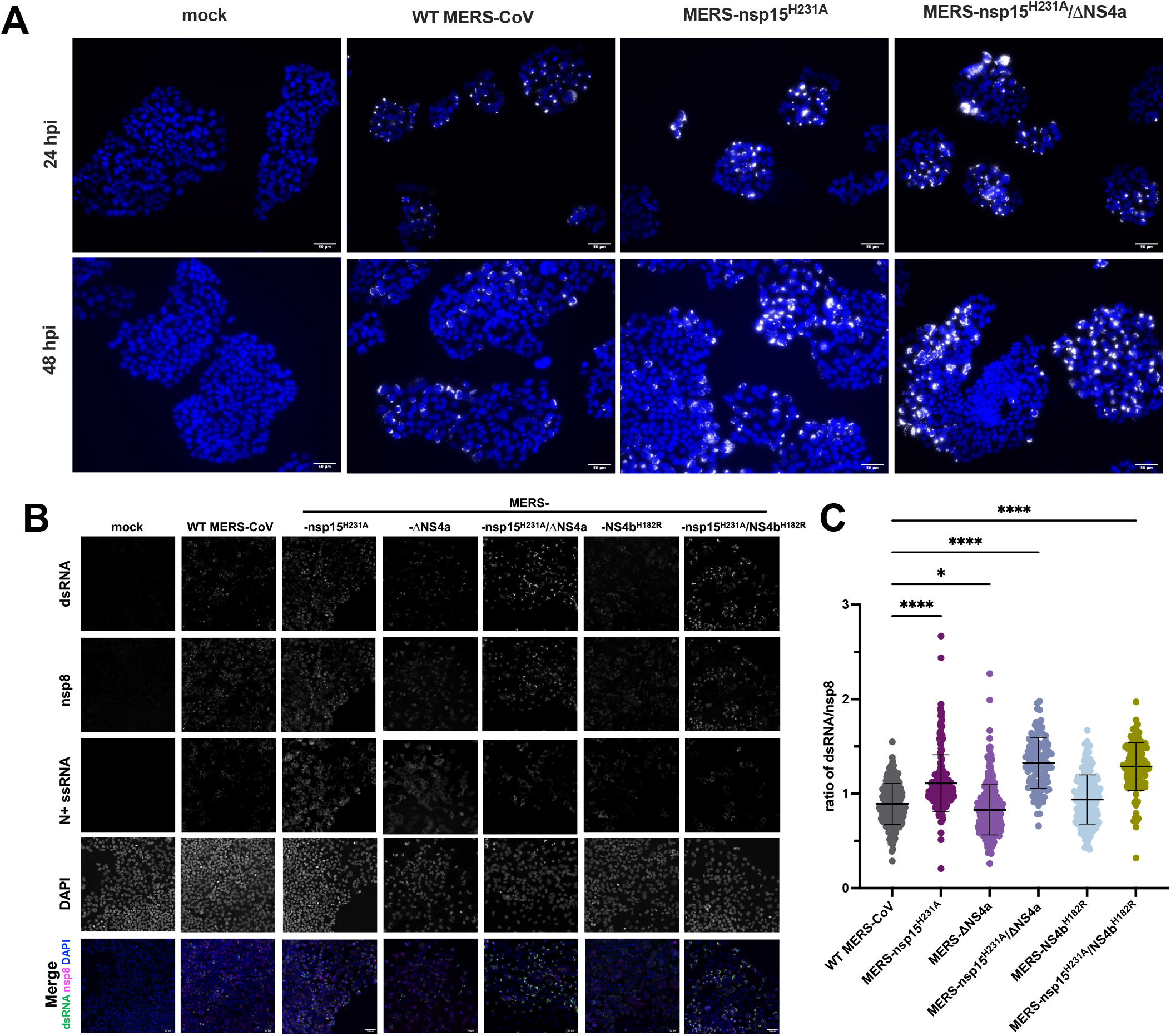
dsRNA expression is increased when EndoU is inactive during MERS-CoV infection. (A) Representative dsRNA staining in MERS-CoV infection: A549^DPP4^ cells were infected at MOI=5 and fixed at 24hpi & 48hpi with 4% paraformaldehyde and subjected to IF to detect dsRNA by 20x widefield microscopy. Nuclei were stained by Hoechst, shown in blue, and dsRNA stained by J2 shown in white. Scale bar is 50μm. (B) Representative images for quantification of dsRNA in A549^DPP4^ cells, mock infected (24hpi) or MERS-CoV infected MOI=5 and fixed 48hpi are shown with all single channels (dsRNA, nsp8, N +ssRNA, DAPI). Merged image shows DAPI (blue), nsp8 (magenta), and dsRNA (green) with scale bar for 50μm. dsRNA and nsp8 were stained by IF using J2 and anti-nsp8 serum, N+ssRNA was stained by FISH, and nuclei were stained by Hoechst (DAPI). (C) dsRNA was quantified by graphing the ratio of dsRNA mean gray value (MGV) over nsp8 MGV in each infected ROI. An ROI was defined by thresholding using the Otsu method for N+ssRNA staining in Fiji. Each dot represents the ratio of dsRNA/nsp8 for that ROI. Five to seven representative fields of widefield microscopy at 20X with 1.5x zoom were analyzed per condition. Data shown are from one representative of two independent experiments and mean plus SD is shown. Statistical significance compared to WT MERS-CoV determined by one-way ANOVA. *, *P*≤0.05; **, *P* ≤ 0.01; ***, *P* ≤ 0.001; ****, *P* ≤ 0.0001. Data that were not statistically significant are not labeled.

### MERS-nsp15^H231A^/ΔNS4a and -nsp15^H231A^/NS4b^H182R^ induce IFN and ISG expression

To investigate if activation of the type I/III IFN pathways was increased during infection in the absence of EndoU activity, we infected A549^DPP4^ cells at MOI=5 with the same panel of MERS-CoV recombinant mutant viruses and collected total intracellular RNA at 24 and 48 hpi (Fig 3). We used quantitative reverse transcription polymerase chain reaction (RT-qPCR) to quantify mRNA expression of select IFN genes (*IFNL1* and *IFNB*) and IFN stimulated genes (ISGs) as compared to mock infected cells. We found minimal expression of IFNs or ISGs over mock infected cells, with infection of any of the viruses at 24hpi. However, at 48hpi expression of *IFNL1* and *IFNB* (over mock infected) remains minimal in WT infected cells and is induced to modest and variable extents in cells infected by the single mutants compared to WT infected cells; in contrast IFN mRNAs as well as ISG mRNAs are significantly induced in double mutant MERS-nsp15^H231A^/ΔNS4a and -nsp15^H231A^/NS4b^H182R^ infected cells compared to WT MERS-CoV infection. Similarly, expression (over mock infected) of select ISG mRNAs, *IFIH1, OAS2, IFIT1*, was also significantly increased in RNA from cells infected with MERS-nsp15^H231A^/ΔNS4a and - nsp15^H231A^/NS4b^H182R^ compared to RNA from WT MERS-CoV infected cells. Additionally, ISG expression was also significantly increased during infection with single mutant MERS-nsp15^H231A^ or -ΔNS4a infection alone, compared to WT-MERS-CoV at 48hpi, although less than in double mutant infected cells.

**Figure 3:**
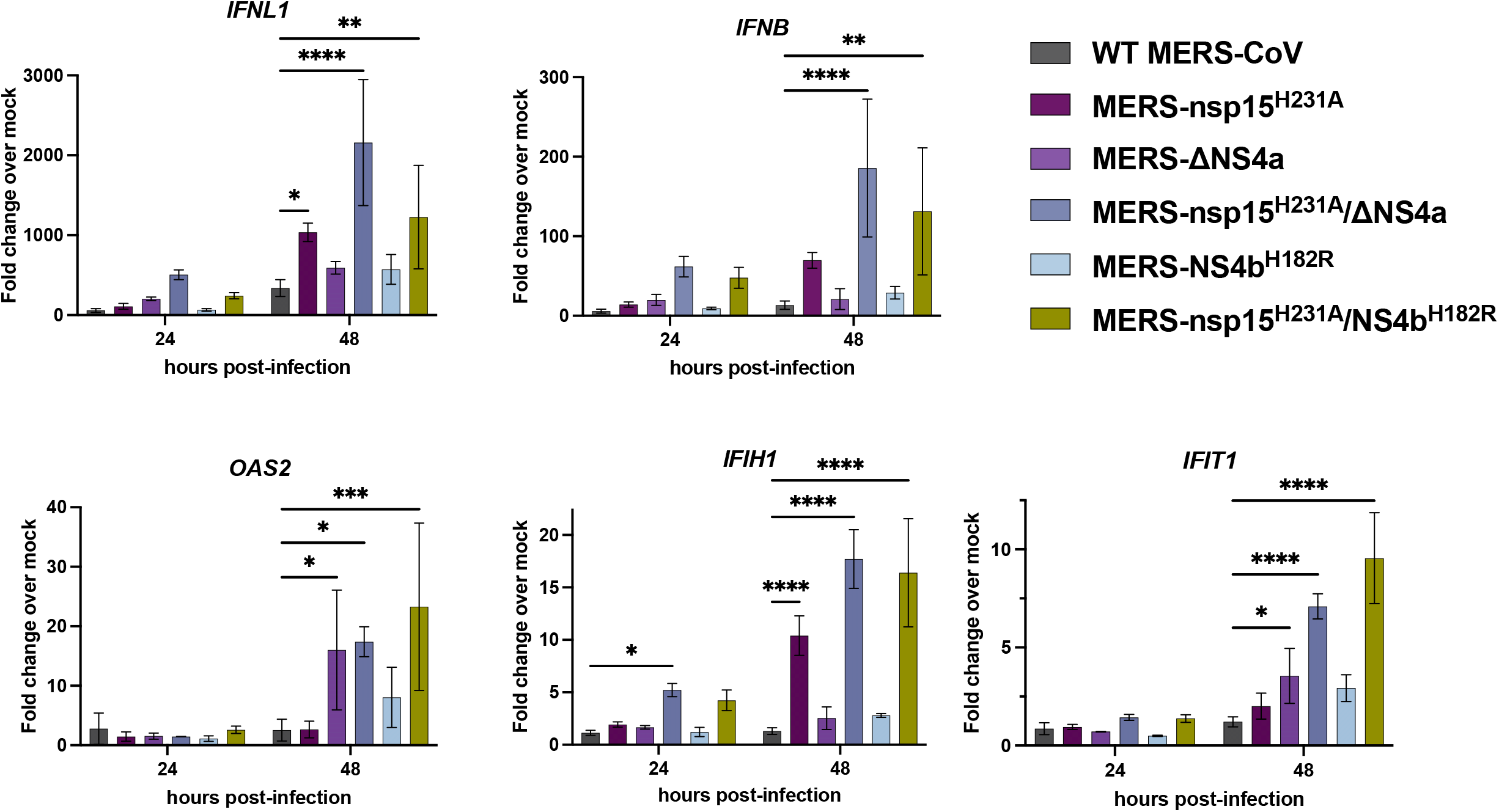
MERS-nsp15^H231A^/ΔNS4a and - nsp15^H231A^/NS4b^H182R^ induce IFN/ISG expression in A549^DPP4^ cells. A549^DPP4^ cells were mock infected or infected in triplicate at an MOI of 5. Total RNA was harvested at 24 and 48hpi and expression of *IFNL1, IFNB, IFIH1, OAS2*, and *IFIT* mRNAs was quantified by qRT-PCR and expressed as fold change over mock infected using the 2^−Δ(Δ*CT*)^ formula. Data are displayed as means ± SD. Data are from one representative of three independent experiments. Statistical significance compared to WT MERS-CoV was calculated two-way ANOVA: *, *P*≤0.05; **, *P* ≤ 0.01; ***, *P* ≤ 0.001; ****, *P* ≤ 0.0001. Data that were not statistically significant are not labeled.

### RNase L and PKR pathways are activated by MERS-CoV double mutants

We next investigated whether RNase L was activated in the absence of EndoU catalytic activity alone or in combination with loss of NS4a expression or NS4b PDE activity. Thus, we infected A549^DPP4^ cells with single and double mutant MERS-CoV as well as WT MERS-CoV at MOI=5, harvested total cellular RNA at 24 and 48hpi, and determined RNase L activation by cleavage of rRNA as indicated by the Agilent RNA Nano 6000 assay (Fig 4A). Sindbis virus (SINV), an unrelated alphavirus that robustly activates RNase L served as a positive control. As we reported previously, RNase L was weakly activated during infection with MERS-NS4b^H182R^ in the absence of PDE activity but not by MERS-ΔNS4a (21, 22). We found that inactivation of EndoU alone (MERS-nsp15^H231A^) also did not lead to strong RNase L activation in A549^DPP4^ cells and that MERS-nsp15^H231A^/ΔNS4a infection weakly activated RNase L as evidenced by slight 28S and 18S rRNA degradation. In contrast, MERS-nsp15^H231A^/NS4b^H182R^ infection, promoted rRNA degradation by 48hpi, indicating more robust RNase L activation in the absence of both EndoU and NS4b PDE enzymatic activities. This contrasts with previous findings in MHV infection of bone marrow derived macrophages where inactivation of either the PDE or EndoU results in robust activation of RNase L and extreme attenuation of replication, to be discussed below.

**Figure 4:**
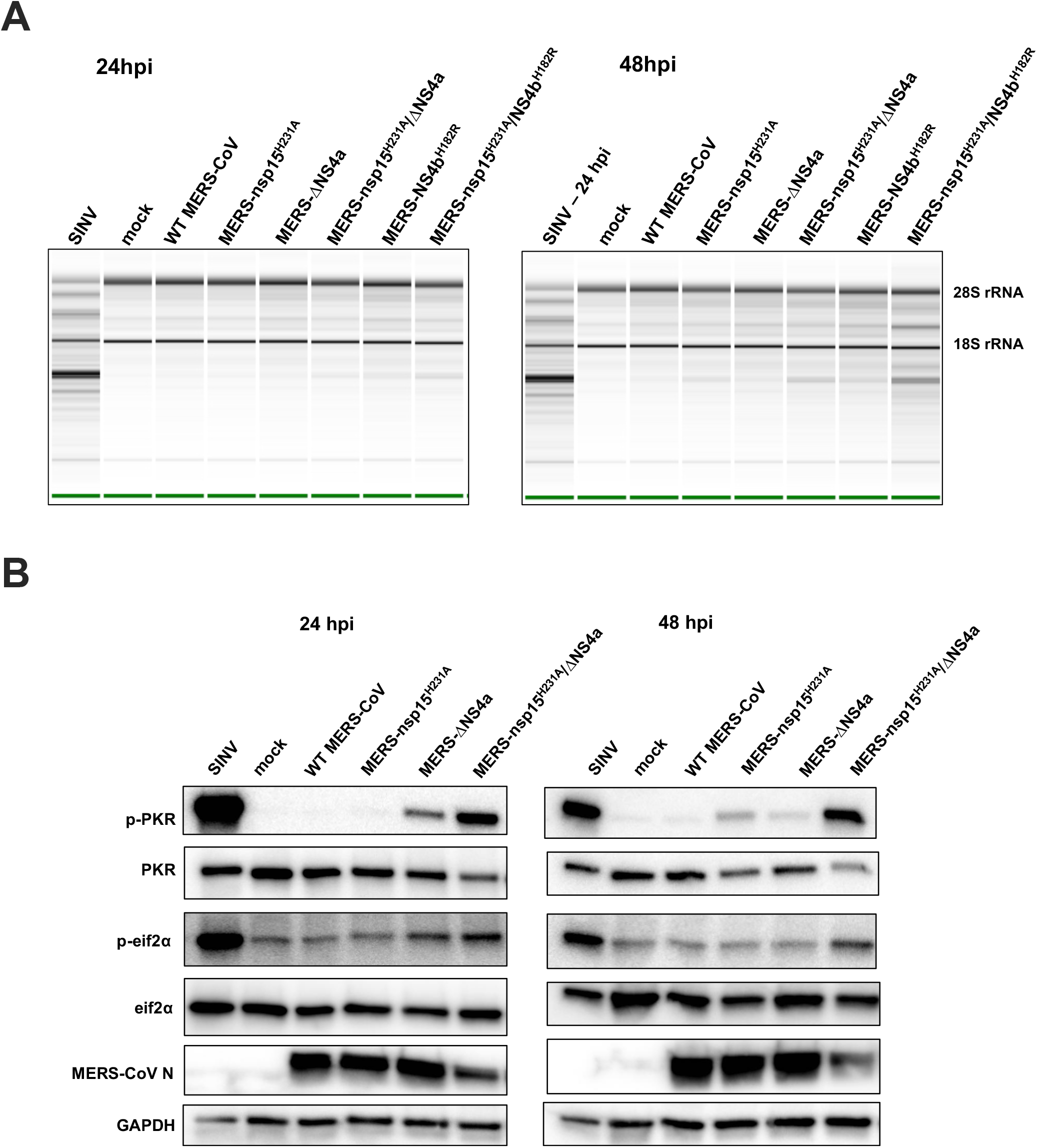
RNase L and PKR are activated in the absence of EndoU activity combined with loss of accessory proteins during MERS-CoV infection. (A) A549^DPP4^ cells were mock infected or infected at MOI=5 and total cellular RNA harvested at 24 and 48hpi. rRNA degradation was assessed on an Agilent Bioanalyzer. 28S and 18S rRNA positions are indicated. Data are from one representative of three independent experiments. (B) A549^DPP4^ cells were mock infected or infected at an MOI of 5 and cell lysates were harvested 24 and 48 hpi. Proteins were separated by SDS-PAGE and immunoblotted with antibodies against phosphorylated PKR (p-PKR), PKR, phosphorylated eIF2α (p-eIF2α), eIF2α, MERS-CoV nucleocapsid (N), and GAPDH. Data are from one representative of four (24 hpi) or two (48hpi) independent experiments.

We also examined induction of the PKR pathway when EndoU is inactivated alone or in combination with loss of NS4a expression. Thus, A549^DPP4^ cells were infected with the single and double MERS-CoV mutants at MOI=5, and protein lysates were harvested at 24 and 48hpi and processed by SDS-PAGE followed by western blotting for detection of PKR activation (Fig 4B). As observed previously, PKR activation was not detected during infection with WT MERS-CoV; however, MERS-ΔNS4a infection induced weak phosphorylation of PKR but not its downstream mediator eIF2α (21, 23, 25). In addition, inactivation of EndoU alone, in infection with MERS-nsp15^H231A^, did not induce strong activation of PKR. Only mild phosphorylation of PKR was observed at 48hpi, late in infection but not at the earlier 24hpi time point, and no phosphorylation of eIF2α was detected. However, during infection with double mutant MERS-nsp15^H231A^/ΔNS4a, strong activation of PKR was observed at both 24 and 48 hpi, indicated by phosphorylation of PKR and eIF2α.

### Knockout of key host immune mediators rescues attenuation observed in MERS-CoV recombinants

Due to replication defects observed in A549^DPP4^ cells and the strong innate immune-stimulatory phenotype observed during MERS-nsp15^H231A^/ΔNS4a and -nsp15^H231A^/NS4b^H182R^ infection, we sought to determine if knock out (KO) of a key mediator in each of the three dsRNA-induced antiviral pathways would rescue replication of these recombinant viruses. We had previously constructed A549^DPP4^ RNase L KO cells (21), and here we additionally generated A549^DPP4^ MAVS KO cells and A549^DPP4^ PKR KO cells to abrogate the host RNase L, IFN, and PKR pathways, respectively. KO of these host immune mediators was confirmed via western blotting (Fig 5A). We further confirmed knockout of the IFN pathway in A549^DPP4^ MAVS KO cells via qPCR in the context of Sendai virus (Cantell strain) infection (39), a virus that induces very high levels of IFNs, showing that both type I (*IFNB*) and type III (*IFNL1*) expression were almost completely eliminated in infected A549^DPP4^ MAVS KO cells compared to WT A549^DPP4^ cells (Fig 5B). We compared replication of WT MERS-CoV with double mutants MERS-nsp15^H231A^/ΔNS4a and -nsp15^H231A^/NS4b^H182R^ in the WT, PKR KO, and MAVS KO A549^DPP4^ cells, along with previously described A549^DPP4^ RNase L KO cells (Fig 5C) (21). Cells were infected at MOI=1 and supernatant samples harvested at 48 hpi. Quantification of infectious virus was completed by plaque assay. In WT A549^DPP4^ cells, we observed a significant attenuation in viral replication in both viral recombinants compared to WT MERS-CoV as expected (Fig 1C). KO of PKR in A549^DPP4^ cells rescued MERS-nsp15^H231A^/ΔNS4a replication to WT MERS-CoV levels, while MERS-nsp15^H231A^/NS4b^H182R^ remained attenuated, consistent with strong activation of the PKR pathway by MERS-nsp15^H231A^/ΔNS4a (Fig 4B). KO of RNase L in A549^DPP4^ cells rescued replication of MERS-nsp15^H231A^/NS4b^H182R^ lacking the PDE activity, while MERS-nsp15^H231A^/ΔNS4a remained attenuated, consistent with our observation of stronger activation of RNase L by MERS-nsp15^H231A^/NS4b^H182R^ (Fig 4A). In MAVS KO cells, replication of both double mutant recombinant viruses (MERS-nsp15^H231A^/ΔNS4a and -nsp15^H231A^/NS4b^H182R^) was rescued to WT MERS-CoV levels. This suggests the particular importance of the IFN pathway in limiting MERS-CoV replication in A549^DPP4^ cells, as elimination of this pathway via MAVS KO was sufficient to restore replication of both recombinant viruses to WT levels.

**Figure 5:**
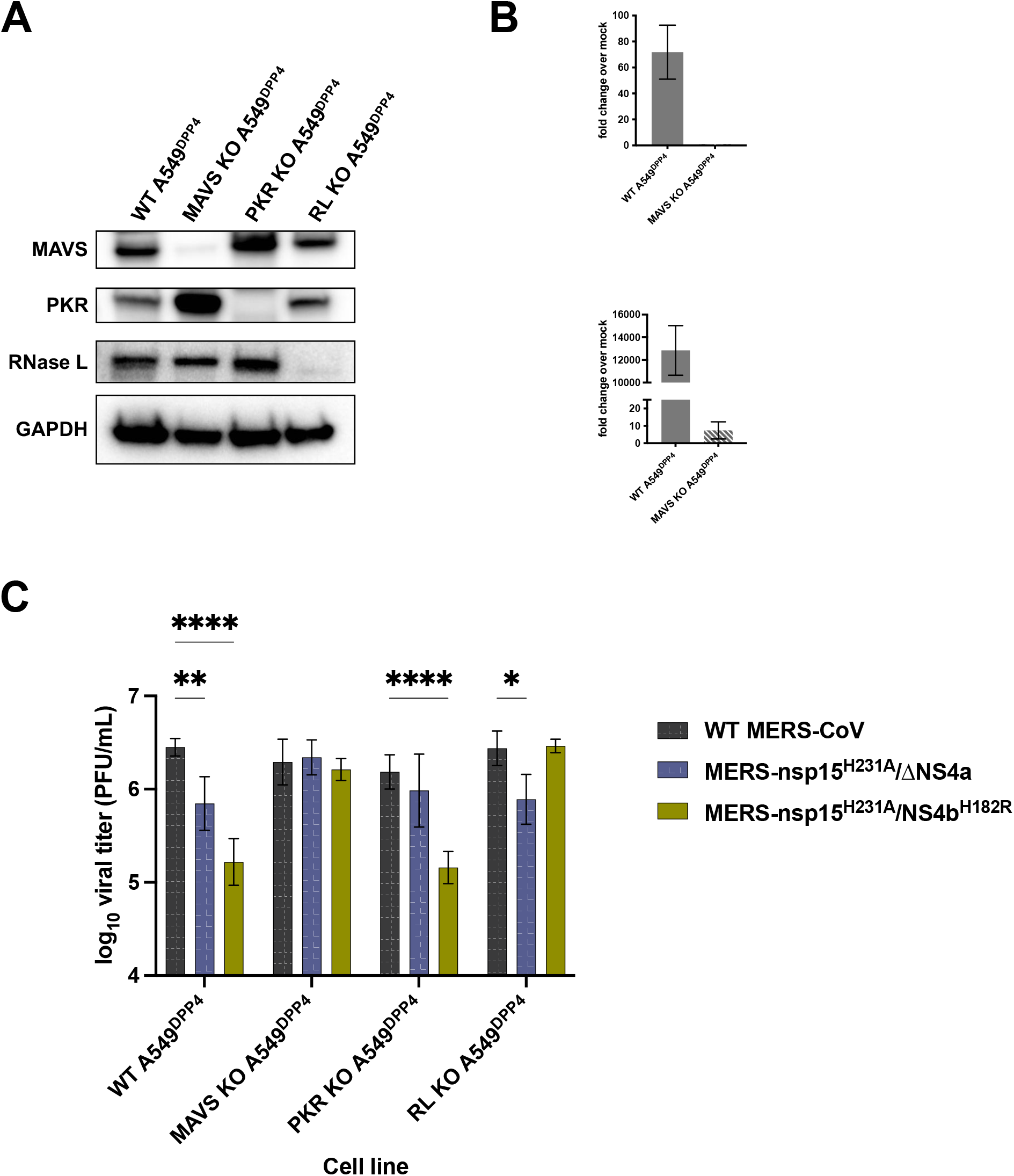
Knockout (KO) of innate immune pathways rescues attenuation in MERS-nsp15^H231A^/ΔNS4a and -nsp15^H231A^/NS4b^H182R^. (A) Cell lysates of WT A549^DPP4^ as well as MAVS KO, PKR KO, and RNase L KO A549^DPP4^ were collected, proteins separated by SDS-PAGE and immunoblotted with antibodies against MAVS, PKR, RNase L, and GADPH. (B) WT A549^DPP4^ and MAVS KO A549^DPP4^ cells were infected with Sendai virus at MOI = 5 and total cellular RNA was harvested at 18hpi. Expression of *IFNL1* and *IFNB* was quantified by RT-qPCR, C_T_ values were normalized to beta-actin and shown as fold-change over mock using the formula 2^−Δ(ΔCT)^. Data are displayed as means ± SD. (C) A549^DPP4^ cell lines: WT MAVS KO, PKR KO, RNase L KO were infected in triplicate at MOI=1 and supernatant samples collected at 48hpi. Viral replication was quantified by plaque assay. Data are displayed as means ± SD. Statistical significance of differences in viral replication for each recombinant virus compared to WT MERS-CoV was calculated by repeated measures two-way ANOVA: *, *P*≤ 0.05; **, *P* ≤ 0.01; ***, *P* ≤ 0.001; ****, *P* ≤ 0.0001. Data that were not statistically significant are not labeled. Data are from one representative of three independent experiments.

### Infection of primary nasal epithelial cell cultures with MERS-CoV recombinants reproduces attenuation and immune-stimulatory phenotype observed in A549^DPP4^ cells

Given the robust activation of dsRNA-induced antiviral innate immune pathways that we observe in A549^DPP4^ cells upon infection with MERS-nsp15^H231A^/ΔNS4a and - nsp15^H231A^/NS4b^H182R^, we sought to compare these viral mutants with WT MERS-CoV in a primary nasal epithelial cell culture system. We grew and differentiated patient-derived nasal samples at an air-liquid interface (ALI) (40) in order to recapitulate the *in vivo* airway epithelium in terms of cell types, including epithelial cells with beating cilia and mucus producing goblet cells, the latter targeted by MERS-CoV (41, 42). We previously showed that these nasal ALI cultures can be productively infected by MERS-CoV (18). To investigate the kinetics of viral replication as well as innate immune induction in these cultures, we infected nasal ALI cultures derived from 2 independent donors with WT MERS-CoV and MERS-nsp15^H231A^/ΔNS4a at MOI = 5. We collected airway surface liquid (ASL) 48, 96, and 144 hpi, and total cellular RNA at 48 and 96 hpi. ASL samples were used to quantify shed virus via plaque assay, and total RNA was used for RT-qPCR analysis for analysis of IFN and ISG expression at each time point. We observed significant reductions in viral replication in MERS-nsp15^H231A^/ΔNS4a compared to WT MERS-CoV in both donors at late time points (Fig 6A), consistent with observations in A549^DPP4^ cells. MERS-nsp15^H231A^/ΔNS4a infection induced significantly more IFN (both *IFNL1* and *IFNB*) and a representative ISG (*IFIT1*) expression compared to WT MERS-CoV in nasal ALI cultures (Fig 6B). We next infected nasal ALI cultures derived from 3 additional donors with WT MERS-CoV, MERS-nsp15^H231A^/ΔNS4a, and -nsp15^H231A^/NS4b^H182R^ to query whether the immune-stimulatory phenotype of both of the double mutants observed in A549^DPP4^ cells could be replicated in this primary epithelial culture system. We observed decreases in viral shedding with both recombinant viruses compared to WT MERS-CoV in 2 of 3 donors at 72hpi (Fig 6C) and hypothesize that the defect in viral replication would be amplified in these cultures at later time points (as observed in Fig 6A). We also observed significantly increased induction of *IFNB* and *IFNL1* mRNA expression in nasal ALIs infected with both recombinant viruses compared to WT MERS-CoV, as well as significantly increased induction of the ISG, *IFIT1* (Fig 6D). We observed variability in IFN responses across donors, likely due to inherent differences between donors in permissiveness to infection and host immune response. Overall, these data demonstrate that the increased innate immune responses observed during infection of A549^DPP4^ cells with both MERS-nsp15^H231A^/ΔNS4a and -nsp15^H231A^/NS4b^H182R^ compared to WT MERS-CoV, as well as the attenuation in these viral recombinants, can be replicated in primary nasal epithelial cultures. This highlights the physiologic relevance of innate immune evasion by MERS-CoV in the upper airway.

**Figure 6:**
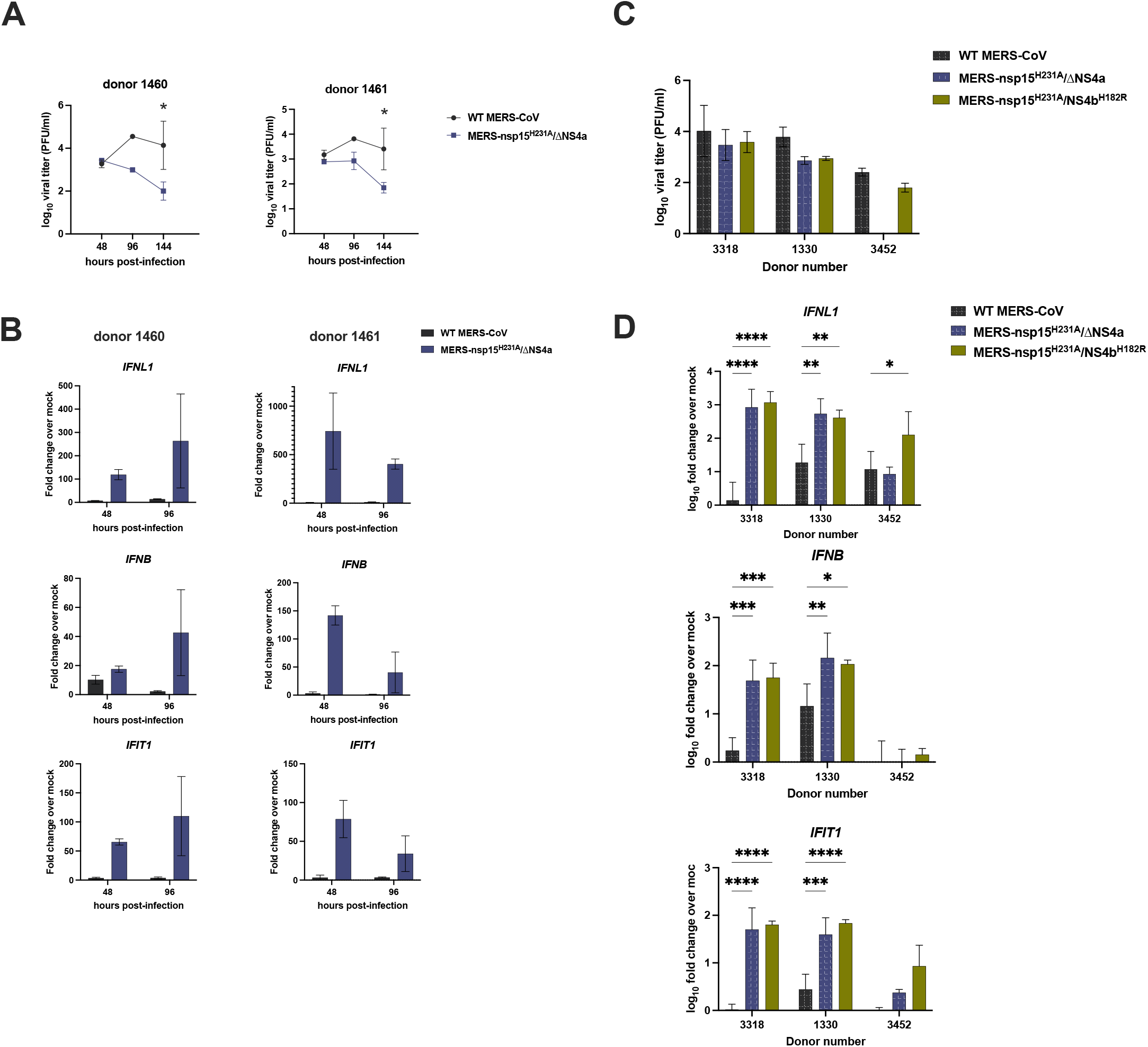
MERS-nsp15^H231A^/ΔNS4a and -nsp15^H231A^/NS4b^H182R^ are attenuated and induce IFN/ISGs in primary nasal epithelial cells. (A) Nasal air-liquid interface (ALI) cultures derived from 2 independent donors were infected in duplicate on the apical cell surface at MOI = 5. Apically released virus was collected at 48, 96, and 144hpi and quantified by plaque assay. (B) After apically released virus was collected from each transwell at the indicated time, total cellular RNA was collected and IFN and ISG mRNA expression was quantified via RT-qPCR. C_T_ values were normalized to 18S rRNA and shown as fold-change over mock using the formula 2^−Δ(ΔCT)^. (C) Nasal ALI cultures from 3 independent donors were infected apically in triplicate at MOI = 5, and apically released virus was collected at 72hpi for quantification by standard viral plaque assay. (D) After collecting apically shed virus, total cellular RNA was collected and IFN and IFIT1 mRNA expression was quantified via RT-qPCR as detailed above using the 2^−Δ(ΔCT)^ method. Data were then log_10_ transformed and displayed as mean ± SD. Statistical significance of differences were calculated by repeated measures two-way ANOVA: *, *P*≤ 0.05; **, *P* ≤ 0.01; ***, *P* ≤ 0.001; ****, *P* ≤ 0.0001. Data that were not statistically significant are not labeled. Data are from one representative of three independent experiments.

## Discussion

The timing of interferon response has been shown to be a crucial factor in determining the outcome of MERS-CoV infection in a mouse model (43). This makes it important to understand how viral antagonist proteins interact with host antiviral pathways to delay or prevent the induction of IFN and related responses. Coronaviruses encode multiple antagonists of host innate immune responses. These include both replicase encoded proteins such as nsp15, conserved among all coronaviruses, and accessory proteins, that are unique for each lineage of coronaviruses, including NS4a and NS4b for MERS-CoV and related merbecoviruses. We have found that MERS-CoV is highly effective at shutting down dsRNA-induced antiviral pathways – IFN signaling, OAS/RNase L and PKR – compared to coronaviruses of other lineages including the sarbecovirus, SARS-CoV-2. Indeed, WT MERS-CoV induces minimal IFN and ISGs, and fails to activate either the OAS/RNase L or PKR pathways, both of which are activated during infection with SARS-CoV-2 (18). We suggest that this is due to expression of the NS4a and NS4b, lineage specific proteins, in combination with the conserved nsp15.

We sought to build on our understanding of the interactions of MERS-CoV accessory proteins NS4a and NS4b with host dsRNA-induced innate immune pathways, by assessing their activities in combination with the nsp15 endoribonuclease. EndoU activity has been shown to limit accumulation of dsRNA and thereby block induction of IFN, OAS/RNase L, and PKR in the context of other coronavirus infections including murine coronavirus MHV, human coronavirus 229E, and porcine epidemic diarrhea virus (PEDV) (33, 34, 44), but the role of EndoU in innate immune evasion during MERS-CoV infection has not before been investigated. Thus, to further understand MERS-CoV antagonism of innate immune pathways and how the effects of EndoU may be combined with those of viral accessory proteins, we constructed a series of recombinant mutant viruses, either with single mutations in antagonist proteins (MERS-nsp15^H231A^, MERS-ΔNS4a, MERS-NS4b^H182R^) or combinations of nsp15^H231A^ with either of the NS4a (MERS-nsp15^H231A^/ΔNS4a) or NS4b (MERS-nsp15^H231A^/ NS4b^H182R^) mutations.

We found that none of the recombinant viruses expressing single mutations exhibited significant replication defects in an immune-competent airway-derived epithelial cell line, A549^DPP4^. This was consistent with our previous findings with a previous set of mutants including NS4b^NLS^ and an NS4ab deletion mutant that lacked expression of both NS4a and NS4b (21). However, we found that combined inactivation of EndoU with ablation of NS4a expression or inactivation of the PDE domain of NS4b in each of the double mutants (MERS-nsp15^H231A^/ΔNS4a and MERS-nsp15^H231A^/ NS4b^H182R^) caused a significant replication defect of approximately one log_10_ PFU/mL in A549^DPP4^ cells (Fig 1). We propose that these MERS-CoV encoded innate immune antagonist proteins (nsp15 (EndoU), NS4a, and NS4b) play overlapping roles by reducing accumulation of dsRNA (EndoU), reducing sensing of dsRNA by PRRs (NS4a), and by directly antagonizing RNase L activity (NS4b PDE), leading to attenuation of replication.

To add to the physiologic relevance of this study, we infected primary cultures of patient derived nasal epithelial cells with WT MERS-CoV and double mutants. The nasal cavity is the primary site encountered by respiratory viruses like MERS-CoV, and these cultures reproduce various features and cell types of the *in vivo* airway (including mucus-producing goblet cells targeted by MERS-CoV, as well as ciliated epithelial cells). Thus, it presents an optimal system in which to study viral immune evasion. We found that MERS-nsp15^H231A^/ΔNS4a and -nsp15^H231A^/NS4b^H182R^ are attenuated for replication and induce the IFN pathway significantly as compared to WT MERS-CoV in this culture system (Fig 6). Thus, MERS-CoV nsp15 EndoU, NS4a, and NS4b play pivotal roles in evading host innate immunity in the upper airways well as in lung-derived cell lines.

Sensing of viral dsRNA is the first step in inducing activation of all three pathways. We observed increased accumulation of dsRNA during MERS-CoV infection when the EndoU catalytic domain was inactivated (MERS-nsp15^H231^, -nsp15^H231A^/ΔNS4a, -nsp15^H231A^/NS4b^H182R^) as detected by immunofluorescence staining with J2 monoclonal antibody. This is similar to reported findings in the MHV system (33, 34). Direct quantification of dsRNA/cell was difficult due to extensive cell-to-cell fusion in MERS-CoV infected cells. Thus, instead we compared the ratios of mean fluorescence intensities of dsRNA to viral protein nsp8 expression, and we observed significant increases in the ratio of dsRNA to nsp8 in MERS-nsp15^H231^, - nsp15^H231A^/ΔNS4a, and -nsp15^H231A^/NS4b^H182R^ compared to WT MERS-CoV in A549^DPP4^ cells (Fig 2).

While EndoU limits accumulation of dsRNA, the accessory proteins are required to more completely suppress the host pathways responding to dsRNA. A mutant with catalytically inactive EndoU (MERS-nsp15^H231A^) induced a modest increase in IFN/ISG mRNA responses. However, a double mutant with inactivated EndoU in combination with either ablation of NS4a expression (MERS-nsp15^H231A^/ΔNS4a) or inactivation of NS4b PDE catalytic activity (MERS-nsp15^H231A^/NS4b^H182R^) significantly increased induction of IFN and ISG mRNA expression late in infection of both lung derived A549 cells and primary nasal cells (Fig 3, 6).

We found that both NS4a and nsp15 EndoU expression are required to fully block activation of PKR. In the absence of NS4a expression, MERS-CoV-ΔNS4a infection promoted only mild activation of PKR (21). However, in cells infected by MERS-nsp15^H231A^/ΔNS4a in the absence of both NS4a and EndoU activity, the PKR pathway was strongly activated at both 24 and 48hpi, evidenced by phosphorylation of PKR and downstream substrate eIF2α (Fig 4B). Interestingly, infection with MERS-nsp15^H231A^ did not promote significant phosphorylation of PKR, indicating that while both NS4a and nsp15 EndoU are required to fully block activation of PKR, NS4a may play a more significant role in blocking induction of this pathway.

As with the PKR pathway, we found that both EndoU and an accessory protein were required to suppress the OAS/RNase L response. Inactivation of either EndoU or the NS4b PDE activity led to only weak activation of RNase L. We observed the most robust activation of RNase L during infection with MERS-nsp15^H231A^/NS4b^H182R^ expressing inactive EndoU as well as inactive PDE. In addition, the binding of NS4a to dsRNA may also contribute to blocking activation of the OAS/RNase L pathway as suggested by observed weak activation of RNase L during infection with MERS-nsp15^H231A^, and somewhat increased with MERS-nsp15^H231A^/ΔNS4a (Fig 4A).

In contrast to the current findings, we reported previously that in the MHV system, inactivation of either the NS4b PDE or EndoU promoted robust RNase L activation (45). The differences observed between MHV and MERS-CoV infection suggest that the PDE and/or EndoU may function differently in the two viral systems. MERS-CoV NS4b is localized mostly in the nucleus due to its NLS, and as such may be less effective as a PDE than the MHV NS2 which lacks an NLS and is completely cytoplasmic. In the MHV system, when expressed in the nucleus, a PDE (AKAP7) was unable to effectively prevent activation of RNase L and rescue virus replication of mutant MHV-NS2^H126R^ with an inactive PDE, but when the same PDE was expressed in the cytoplasm it prevented activation of RNase L and restored replication of MHV-NS2^H126R^ to WT MHV levels (46). Additionally, our MHV studies were carried out in bone marrow derived macrophages which express very high levels of OAS genes (17, 47) which may lead to increased activation of RNase L, and both antagonists may be necessary to prevent RNase L activation in a cell type dependent manner. These findings highlight the importance of studying the function of these proteins in the context of different coronavirus infections and different cell types.

To verify that innate immune activation was restricting each of the double mutants, we constructed A549^DPP4^ cell lines with host innate immune mediators in each of the three dsRNA-induced antiviral pathways knocked out. We found that abrogation of specific immune pathways can rescue viral attenuation of MERS-nsp15^H231A^/ΔNS4a and -nsp15^H231A^/NS4b^H182R^ to WT MERS-CoV levels. While PKR KO rescued only MERS-nsp15^H231A^/ΔNS4a replication, RNase L KO rescued only MERS-nsp15^H231A^/Ns4b^H182R^, which was consistent with these recombinant viruses robustly activating each of these pathways. These data support the contributions of NS4a and NS4b to suppressing the PKR and OAS/RNase L pathways, respectively. Interestingly, replication by both of these recombinant viruses was rescued to WT MERS-CoV levels in MAVS KO A549^DPP4^ cells, suggesting that the IFN pathway is particularly important in limiting MERS-CoV replication when activated. This may also be explained in part by the fact that PKR and OASs are ISGs themselves. These observations underscore the importance of antagonism of these pathways by MERS-CoV, and provide further evidence that these host immune pathways play pivotal roles in limiting CoV replication in the airway.

There are many unanswered questions about interactions of coronaviruses with host innate immunity during infection by highly pathogenic coronaviruses. Indeed, each lineage of coronavirus encodes different groups of accessory proteins and interacts differently with host innate pathways which will confer different interactions with these pathways. For example, SARS-CoV-2 accessory proteins do not include homologs of NS4a or NS4b and unlike MERS-CoV, SARS-CoV-2 activates the dsRNA-induced innate immune pathways (IFN, PKR, OAS/RNase L) and WT SARS-CoV-2 is restricted in replication by RNase L (18). We hypothesize that loss of EndoU activity during SARS-CoV-2 infection may lead to more robust activation of these pathways and detrimental effects on replication and pathogenesis, currently under investigation. Finally, EndoU activity of nsp15 is an attractive antiviral drug target for current and future highly pathogenic coronaviruses, as it is highly conserved in coronaviruses.

## Materials and Methods

### Recombinant viruses

Recombinant MERS-CoV viruses were made using lambda red recombination with the MERS-CoV Bacterial Artificial Chromosome (BAC) as previously described (37, 48, 49). The following primers were made to generate the mutations (bolded) to create the recombinant viruses:

MERS-**nsp15^H231A^**:

F 5’-GTGATGTTTTCATTAAGAAGTATGGCTTGGAAAACTATGCTTTTGAG**GC**CGTAGTCTATGGA GACTT-3’

R 5’-CGCCTAACGTAGTATGAGAGAAGTCTCCATAGACTACG**GC**CTCAAAAGCATAGTTTTCCA-3’ MERS-**ΔNS4a**: (amino acids 11 and 12 replaced by stop codons)

F 5’-GAACTCTATGGATTACGTGTCTCTGCTTAATCAAATTTG**AT**AGAAGTACCTTAACTCACC-3’

R 5’-TGTACAAACAAGTAGTATACGGTGAGTTAAGGTACTTCT**AT**CAAATTTGATTAAGCAGAG-3’ MERS-**NS4b^H182R^**:

F 5’-GTTCAGGGATTTTCCCTTTACCATAGTGGCCTCCCTTTAC**G**TATGTCAATCTCTAAATTG-3’

R 5’-GTAACATCATCCAGTGCATGCAATTTAGAGATTGACATA**C**GTAAAGGGAGGCCACTATGG-3’

MERS-**nsp15^H231A^/ΔNS4a**: the same mutations as above combined on one BAC

MERS-**nsp15^H231A^/ NS4b^H182R^**: the same mutations as above combined on one BAC

All amplified KanI using the following additions onto the above primers:

F 5’-AGGATGACGACGATAAGTAGGG-3’ R 5’-GCCAGTGTTACAACCAATTAACC-3’

Passage 0 (P0) viruses were recovered by Invitrogen™ Lipofectamine™ 2000 transfection reagent on African Green Monkey VeroCCL81 cells using 1.25ug of BAC DNA. Cells were monitored for cytopathic effect and cells and supernatant harvested 5-7 days after transfection. P0 virus was freeze thawed, cells removed by centrifugation and passaged onto Vero CCL81 cells to generate P1 stock. A P2 stock was generated by using low MOI (0.05) of P1 stock and infecting Vero CCL81 cells, freeze thawed to harvest, cells removed and supernatant used as the stock virus for experiments. Quantification of infectious virus was done by standard viral plaque assay on Vero CCL81 cells (18).

Sindbis virus Girdwood (G100) (SINV) was obtained from Dr. Mark Heise, University of North Carolina, Chapel Hill, and prepared as previously described (50). Sendai virus was obtained from Dr. Carolina Lopez, Washington University.

### Cell lines

Vero CCL81 cells were cultured in DMEM+10% FBS, sodium pyruvate, and HEPES. Human A549^DPP4^ cells were cultured in RPMI 1640 supplemented with 10% FBS and penicillin-streptomycin. A549^DPP4^ and A549^DPP4^ *RNASEL* knock out (KO) cells were previously described (21). A549^DPP4^ *PKR* KO cells were constructed using the same Lenti-CRISPR system and guide RNA sequences as previously described (11, 51). A549^DPP4^ *MAVS* KO cells were constructed using the same Lenti-CRISPR system as previously described (51) but with the following guide RNAs: sgMAVS-2 forward: 5’-CACCGGGGTCTCCTGGACAGGCATG-3’ and sgMAVS-2 reverse: 5’-AAACCATGCCTGTCCAGGAGACCCC-3’

### Nasal air-liquid interface (ALI) cultures

Primary epithelial cultures were derived from human nasal mucosal specimens acquired from residual clinical materials obtained during surgery performed through the Department of Otorhinolaryngology (Head and Neck Surgery) at the University of Pennsylvania after informed consent was obtained. Patients with history of systemic disease or on immunosuppressive medications were excluded as previously described (32). Acquisition and use of these nasal specimens have been approved by the University of Pennsylvania Institutional Review Board (protocol #800614). Cultures were grown and differentiated on transwell inserts as previously described (32). Once fully differentiated (approximately 6 weeks post-seeding), basal media was replaced and the cultures were washed apically with PBS three times prior to infection apically with MERS-CoV.

### MERS-CoV infections and titration

Viruses were diluted in serum-free DMEM and added to cells for adsorption for 1 hour at 37°C. Cells were washed three times with PBS (for growth curves and RNA, no washes for protein or immunofluorescence samples) and fed with RPMI+2% FBS for A549^DPP4^ or DMEM+2%FBS for Vero CCL81. 150 μl of supernatant was collected at the times indicated and stored at −80°C for titration by plaque assay on Vero CCL81 cells as previously described (18). All infections and virus manipulations were conducted in a biosafety level 3 (BSL3) laboratory using appropriate personal protective equipment and protocols.

### Immunofluorescence (IF) and fluorescent *in situ* hybridization (FISH) staining

At indicated times post-infection cells were fixed with 4% paraformaldehyde for 30 minutes at room temperature. Cells were then washed three times with PBS and then dehydrated with increasing amounts of ethanol: 50%, 70%, and 100% for 5 minutes each. Samples were stored at 100% ethanol at −20°C overnight. Cells were rehydrated with 70% then 50% ethanol for 2 minutes each, washed with PBS once and then permeabilized for 10 minutes with PBS+0.1% Triton-X100. Cells were then blocked in PBS+0.1% Triton-X100 and 2% BSA containing RNaseOUT™ (Invitrogen™ Cat #10777019) for 30 minutes at room temperature. Primary antibodies were diluted in block buffer and incubated at room temperature for one hour. Cells were washed three times with PBS and then incubated at room temperature for 30 minutes with secondary antibodies diluted in block buffer. All steps following secondary antibody addition were carried out in the dark. Cells were washed three times with PBS. For IF only, nuclei were stained with DAPI diluted in PBS for 5 minutes at room temperature, and coverslips were mounted onto slides using Invitrogen™ ProLong™ Diamond Antifade Mounting reagent for analysis by widefield microscopy. For combined IF/FISH staining, cells were washed three times with PBS (as stated above) and then fixed again with 4% paraformaldehyde for 10 minutes at room temperature. Cells were washed with 2X SSC for 5 minutes and then 5 minutes with FISH wash buffer (10% formaldehyde in 2x SSC). FISH probes complementary to the N gene of MERS-CoV (see table below) conjugated to CAL Fluor Red 610 (Stellaris custom order) were diluted 1:50 in nuclease free water.

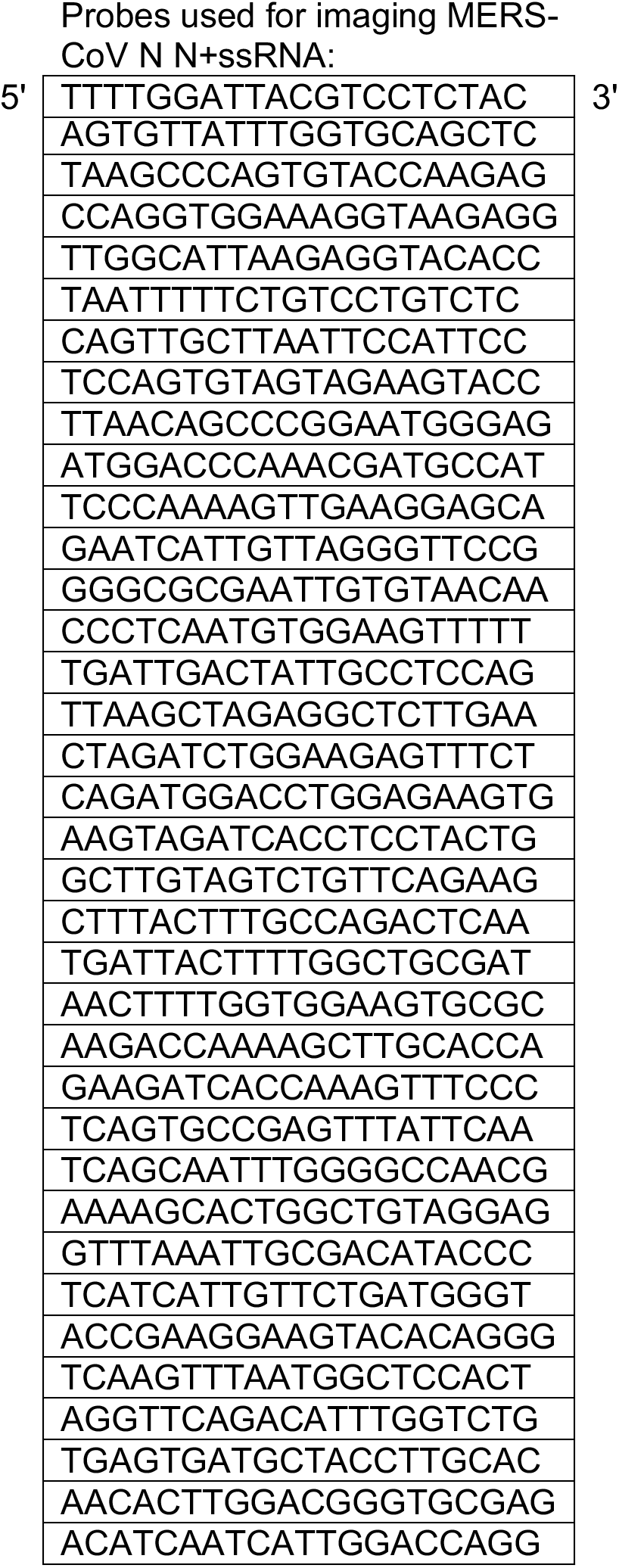

Probes were diluted 1:50 in hybridization buffer (10% w/v dextran sulfate in 10% formaldehyde in 2x SSC) and incubated with samples overnight at 37°C in a humidity chamber. Samples were washed with FISH wash buffer for 30 minutes at 37°C, nuclei stained with DAPI diluted in FISH wash for 5 minutes at room temperature, and then cells were washed for 5 minutes with 2X SSC. Coverslips were mounted onto slides as stated above. DsRNA was detected using commercial monoclonal antibody J2 (Scions) at 1:500 and nsp8 using anti-nsp8 rabbit serum 1:500 (obtained from Dr. Mark Denison, Vanderbilt University). Secondary antibodies were diluted 1:1000, all highly cross-adsorbed IgG (H+L) from Invitrogen: goat anti-mouse AF488 (Cat #: A11029) and goat anti-rabbit AF647 (Cat #: A32733).

Widefield microscopy was done using either Nikon Eclipse Ti2 using a Nikon 20x objective and NikonDS-Qi1Mc-U3 12-bit camera or Nikon Eclipse Ti2 using a Nikon 20x objective and Hamamatsu digital camera C13440. Fiji was used for quantification of dsRNA: N +ssRNA staining was used to generate a mask by setting a threshold using the Otsu method. These thresholded images were as a mask to mark infected regions of interest (ROI). These ROIs, due to syncytia formation, marked infected areas of cells rather than singular cells. The mean gray value (MGV) of fluorescence signal of dsRNA or nsp8 was measured within each ROI across several images per infection condition. Five to seven fields of view at 20x with 1.5x zoom were analyzed for each condition. The ratio of MGV of dsRNA over MGV of nsp8 was recorded for each individual ROI and plotted using GraphPad Prism software.

### Western blotting

Cells were washed once with ice-cold PBS and lysates harvested at indicated times post infection with lysis buffer (1% NP40, 2mM EDTA, 10% glycerol, 150mM NaCl, 50mM Tris HCl) supplemented with protease inhibitors (Roche – cOmplete mini EDTA-free protease inhibitor) and phosphatase inhibitors (Roche – PhosStop easy pack). After 5 minutes lysates were harvested, incubated on ice for 20 minutes, centrifuged for 20 minutes at 4°C and supernatants mixed 3:1 with 4x Laemmli sample buffer. Samples were heated at 95°C for 5 minutes, then separated on 4-15% SDS-PAGE, and transferred to polyvinylidene difluoride (PVDF) membranes. Blots were blocked with 5% nonfat milk or 5% BSA in TBST and probed with the following antibodies diluted in the same block buffer:

| Primary Antibody | Antibody species | Blocking buffer | Dilution | Catalog number |
| --- | --- | --- | --- | --- |
| p-PKR (phospho-T446) [E120] | rabbit | 5% milk/TBST | 1:1000 | Abcam 32036 |
| PKR (D7F7) | rabbit | 5% milk/TBST | 1:1000 | Cell Signaling Technology 12297S |
| peif2α (S51) | rabbit | 5% BSA/TBST | 1:1000 | Cell Signaling Technology 9721S |
| eif2α | rabbit | 5% BSA/TBST | 1:1000 | Cell Signaling Technology 9722S |
| MERS-CoV N | mouse | 5% milk/TBST | 1:2000 | 40068-MM10 (Sino Biological) |
| GAPDH (14C10) | rabbit | 5% milk/TBST | 1:2000 | Cell Signaling Technology 2118S |
| MERS-CoV NS4a | rabbit | 5% milk/TBST | 1:500 | obtained from Dr. Luis Enjuanes, Spanish National Centre for Biotechnology |
| MERS-CoV NS4b | rabbit | 5% milk/TBST | 1:500 | obtained from Robert Silverman, Cleveland Clinic |
| MAVS (E-3) | mouse | 5% milk/TBST | 1:250 | Santa Cruz-166583<br>*Used secondary antibody: goat anti-mouse IgG-HRP Santa Cruz-2005 |
| RNase L | mouse | 5% milk/TBST | 1:1000 | obtained from Dr. Robert Silverman, Cleveland Clinic |
| Beta-tubulin | rabbit HRP-linked | 5% milk/TBST | 1:5000 | Abcam 21058 |
| **Secondary Antibody** |  |  |  |  |
| goat anti-rabbit IgG | HRP linked | same as primary | 1:3000 | Cell Signaling Technology 7074S |
| goat anti-mouse IgG | HRP linked | same as primary | 1:3000 | Cell Signaling Technology 7076S |

Blots were visualized using Thermo Scientific SuperSignal west chemiluminescent substrates (Cat #:34095 or 34080). Blots were probed sequentially with antibodies and in between antibody treatments stripped using Thermo Scientific Restore western blot stripping buffer for 1 hour at room temperature (Cat #: 21059).

### Quantitative PCR (qRT-PCR)

At indicated times post-infection cells were lysed with buffer RLT Plus (Qiagen RNeasy Plus #74136) and RNA extracted following the prescribed protocol. RNA was reverse transcribed into cDNA with a High Capacity cDNA Reverse Transcriptase Kit (Applied Biosystems). cDNA was amplified using specific qRT-PCR primers (see Table 3.3), iQ™ SYBR® Green Supermix (Bio-Rad), and the QuantStudio™ 3 PCR system (Thermo Fisher). Fold changes in mRNA level compared to mock infected samples were calculated using the formula 2^−Δ(ΔCt)^(ΔCt = Ct_gene of interest_ – Ct_18S_) and expressed as fold infected/mock-infected. Primer sequences are as follows:

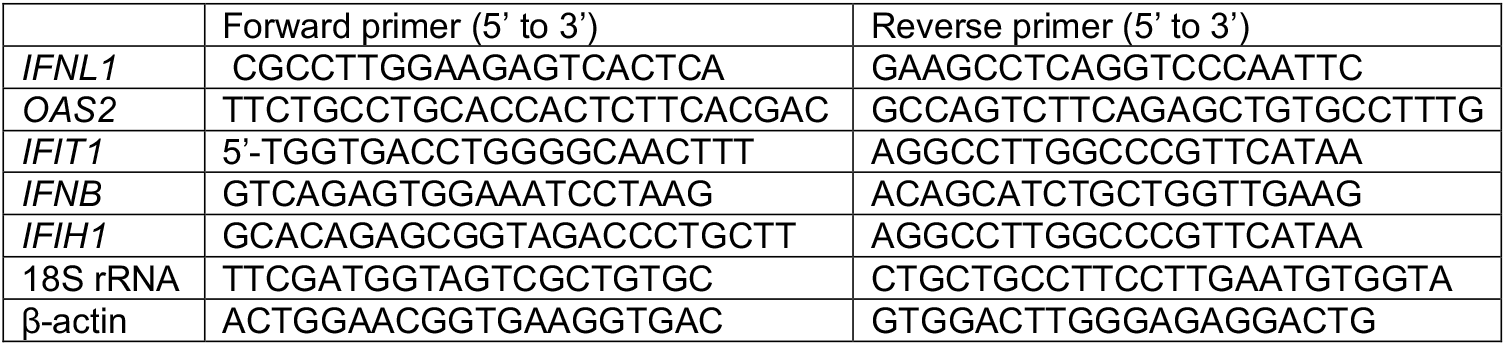

### Analyses of RNase L-mediated rRNA degradation

RNA was harvested with buffer RLT plus (Qiagen RNeasy Plus #74136) and analyzed on an RNA chip with an Agilent Bioanalyzer using the Agilent RNA 6000 Nano Kit and its prescribed protocol (Cat #: 5067-1511).

### Statistical analysis

Plotting of data and statistical analysis were performed using GraphPad Prism software (GraphPad Software, Inc., CA). Statistical significance was determined by comparing mutant viruses to WT MERS-CoV using repeated measures two-way ANOVA for viral replication curves and qRT-PCR and by one-way ANOVA for dsRNA quantification. Displayed significance is determined by p-value (P), where * = P < 0.05; ** = P < 0.01; *** = P < 0.001; **** = P < 0.0001; ns = not significant and in some figures, ns is not displayed on the graph.

## Acknowledgements

We thank the members of the Weiss lab for feedback and discussion of this project, Joshua Hatterschide for assistance with microscopy image analysis, Rudragouda Channappanavar (Oklahoma State University) for construction of MERS-nsp15^H231A^, Nicholas Parenti for performing the Bioanalyzer assays. Mark Heise (University of North Carolina, Chapel Hill) for Sindbis virus and Carolina Lopez (Washington University, St Louis) for Sendai virus. This work was supported by National Institutes of Health grant R01-AI140442 and the Penn Center for Research on Coronaviruses and Other Emerging Pathogens. DMR was supported in part by T32 AI055400 and CEC in part by T32 NS007180.

